# Modeling the evolutionary history of nonclassical monocytes in mammals

**DOI:** 10.1101/2022.10.12.511958

**Authors:** Daniel J. Araujo, Diego Fernandez, Ahmad Alimadadi, Axel Zagal-Norman, Catherine C. Hedrick

**Author notes:** **Cite as:** Araujo DJ, Fernandez D, Alimadadi A, Zagal-Norman A, and Hedrick CC. Modeling the evolutionary history of nonclassical monocytes in mammals. *Under preparation*.

## Abstract

The immune system has experienced major changes in both its organization and function during the evolution of tetrapod animals. Still, the ancestry of specific cell types and the historical relationships between modern immune cells has received little attention. While all tetrapods possess mononuclear blood cells and monocytes, vasculature-patrolling nonclassical monocytes (CD16^+^CD14^-^ in humans, Ly6C^-^ in mice) have so far only been described in mammals. The question of whether nonclassical monocytes are specific to the mammalian lineage has not been answered. Cell types can be described as traits that persist via the inheritance of core regulatory complexes of transcription factors. We utilized transcriptional network analyses on a human monocyte single-cell RNA-sequencing dataset to highlight the core regulatory complex associated with nonclassical monocyte production (nCoRC). We then applied BLAST-based analyses to quantify the conservation of human nCoRC members amongst mammals and nonmammals. We determined that the average sequence similarity of nCoRC members is highest amongst mammals and that such conservation is not observed in non-mammalian tetrapods. We also discovered that this sequence similarity is specifically driven by boreoeutherian placental mammals. Finally, we found the receptor proteins upstream of nonclassical monocyte assembly are also more homologous in mammals than non-mammals. This work provides an evolutionary model in which the capacity for nonclassical monocyte production is unique to the mammalian lineage.

## INTRODUCTION

Tetrapods represent a 350-million-year-old monophyletic group that includes all four-limbed vertebrates and their descendants. The immune system has undergone pronounced architectural changes during the evolutionary history of tetrapods. These alterations include the emergence of adaptive immunity^1^, tissue specialization amongst macrophages^2^, and increases in the inflammatory and phagocytic capacity of peripheral neutrophils^3^. Studies into evolutionary immunology could hence offer blueprints for designing and promoting immune cells with certain therapeutic properties. Nevertheless, determining the ancestry of these cell types remains challenging due to the mutability of immune cells *in vivo* and the paucity of fossilized cellular material^4^. Cell types can be defined as evolutionary units that persist via the inheritance of core regulatory complexes (CoRCs) of transcription factors^5,6^. CoRCs allow cells to move along paths of ontogeny by inducing certain transcriptional states in response to particular signals. Thus, CoRCs permit cell types to enact specific physiological modules and provide testable hypotheses of cellular ancestry.

Monocytes are myeloid cells that act as innate immune responders and which result from the action of a transcriptional circuit involving IRF8 and KLF4 within common monocyte precursors^7–9^. Classical monocytes (CD14+CD16^-^ in humans, Ly6C^hi^CCR2^hi^ in mice) give rise to a diverse pool of cells known as intermediate monocytes (CD14^+^CD16^+^ in humans, Ly6C^hi^Treml4^+^ in mice)^10–12^. Intermediate monocytes in turn form a transcriptional continuum leading to nonclassical monocytes (CD14^-^CD16^+^ in humans, Ly6C^lo^CCR2^lo^ in mice)^7,11,12^. Conversion of classical monocytes into nonclassical monocytes is controlled by the transcription factors C/EBPβ, KLF2, and NR4A1^11,13^. Nonclassical monocytes crawl along the peripheral vasculature, scavenge endothelial debris, and protect against atherosclerotic damage to the endothelium^14–17^. Production of these “patrolling” monocytes is dependent on DLL1^+^ and CSF1^+^ sinusoidal niches in bone marrow^18,19^. While these cells are likely produced in the peripheral circulation via the activity of C/EBPb^11^, the signals controlling this process are not fully understood.

In addition to humans and mice, nonclassical monocytes are present in the blood of rats^20^, dogs^21,22^, cows^23^, sheep^24^, bats^25^, horses^26^, and macaques^27^. These species all fall inside the clade of *Boreoeutheria*, which denotes one of the two major divisions of placental mammals^28^. All extant tetrapods possess mononuclear blood cells and monocytes^3,29^, but whether nonclassical monocytes are specific to mammals or restricted to a subset of mammals has not been determined. Given the conserved features of the mammalian cardiovascular system^30,31^ and the intimate link between nonclassical monocytes and vascular signals in humans and mice, we hypothesized that nonclassical monocytes are a synapomorphy of the mammalian lineage. To test this hypothesis, we analyzed a scRNA-seq dataset derived from healthy human PBMCs to define the CoRC of nonclassical monocytes. We then employed comparative gene homology to show that the members of the nonclassical monocyte CoRC are highly conserved amongst boreoeutherian mammals when compared to transcription factors associated with other cell types. Our systematic approach revealed that this homology is not shared by marsupials, monotremes, or non-mammalian tetrapods.

## METHODS

### Systematic selection of species

We chose 69 species of tetrapods (including humans) with annotated genomes for our analysis. Of these organisms, 67 species were amniotes and 2 were amphibians. The amniotes included 7 lepidosaurs, 6 testudines, 16 archosaurs, and 38 mammals. We excluded non-tetrapod vertebrates (lungfish, lobe-finned fishes, ray-finned fish, cartilaginous fish, etc.) because of the additional genomic duplications^32^ and unique genomic features in these lineages^33,34^. Phylogenetic trees of the relationships between tetrapods were retrieved from TimeTree^35,36^. Descriptive information on all the species we selected is presented in **Supplementary Table 1**.

### SCENIC analysis

We focused on the publicly available scRNA-seq dataset from Watanabe et al., 2020^37^ for our transcriptional network analysis. Original FASTQ files deposited in the NCBI Gene Expression Omnibus (GEO) database (GSE142444) were downloaded from Sequence Read Archive (SRA) repository. The quality of sequencing reads was investigated using the FastQC software^38^. The CellRanger v7.0 package was used to align the reads to the human reference genome (GRCh38) and to quantify expression^39^.

The Seurat v4.0 R package^40^ was utilized to perform analysis on the raw count matrices from the CellRanger process. To exclude potential droplets and doublets, cells with less than 250 and more than 2500 features, respectively, were excluded. Cells with more than 7.5% of counts originating from mitochondrial reads were also considered low-quality/dying cells and excluded from our analysis. Counts were normalized using the LogNormalize method with scale.factor of 10000. The top 2000 variable genes were used to perform PCA analysis. Since every patient sample was sequenced during a different run, both cell clustering and UMAP generation were performed using the top 30 Harmony dimensions calculated with the top 30 PC dimensions^41^. Cell populations were identified using canonical markers and gene signatures provided in GSE25913.

Using the normalized gene expression values as input, gene regulatory network analysis was implemented by SCENIC v.1.3.1 in R to identify transcription factor regulons^42,43^.

### Comparing protein sequence similarity between species

#### Retrieval of reference gene sequences

Human genes served as a baseline for comparing sequence similarity scores against orthologous proteins of different species. In brief, gene sequences were first retrieved from the NCBI database and only sequences included in the MANE project were selected for downstream analyses. Isoforms and duplicated entries were resolved by selecting those gene IDs with the highest quality of annotations and completeness.

#### Comparing protein sequence similarity between species

Human protein sequences for each gene were then BLASTed against the corresponding protein products from the Ensembl reference genomes of the other 68 tetrapods included in this study (*release 106 - accessed March 2022*). Reverse searches of every retrieved hit against the human database were performed. Homology was scored by selecting those proteins that corresponded to bidirectional best hits between the queried organism and humans. Bidirectional best hits were defined as pairs of proteins that were more similar to each other than either was to any other corresponding gene in the opposing genome. When genes did not produce a bidirectional best hit, we selected the match with the highest bit-score.

Sequence similarity scores were reported as the percentage identity between two orthologs as indicated by the BLOSUM62 substitution scoring matrix. While we included the sequence similarity scores of every species to obtain clade averages, all data are reported with a 40% cutoff where possible, as this is a traditional threshold for inferring homology^44^.

### Visualization and statistical software

GraphPad Prism and R studio were used to plot the data and perform statistical analyses.

## RESULTS

### Nonclassical monocytes are observed in boreoeutherian mammals

Extant members of *Tetrapoda* are divided into amphibians, testudines (turtles), archosaurs (birds and crocodilians), lepidosaurs (lizards and snakes), and mammals (**Figure 1**). Mammals are subsequently divided into three major groups: monotremes, marsupials, and placentals. Monotremes are the most basal mammalian form in that they lay eggs, whereas marsupials produce altricial offspring that require pouches for rearing and placental mammals produce precocial offspring. Placental mammals segregate into the phylogenetic clades *Boreoeutheria* and *Atlantogenata*, with modern boreoeutherian mammals embodying the more diverse family^28^. Nonclassical monocytes are found in the peripheral blood of humans, macaques, rats, mice, bats, dogs, cows, sheep, and horses, all of which are members of *Boreoeutheria* (**Figure 1**). However, little effort has been devoted to identifying orthologous classical, intermediate, and nonclassical monocyte subtypes in other mammalian families or non-mammalian species. It is therefore unknown whether nonclassical monocytes are an inherited trait specific to the boreoeutherian lineage or if they are common to other mammalian families or even non-mammalian tetrapods.

**Figure 1.**
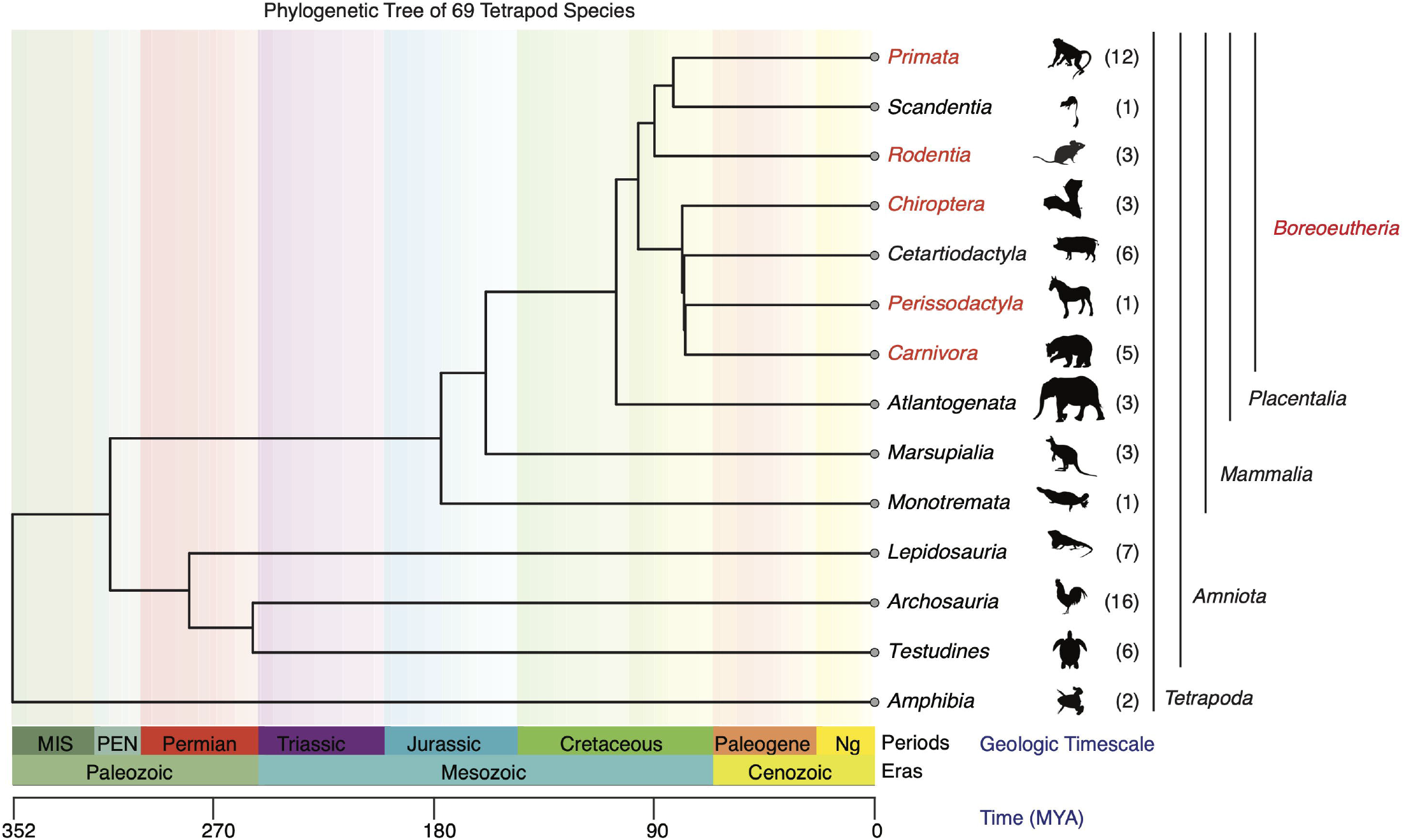
Nonclassical monocytes are exclusive to boreoeutherian mammals. Phylogenetic tree of the consensus relationships between the 69 tetrapod species considered in this study. Numbers to the right of the representative clade icons indicate the count of individual species used in our downstream analyses. Mammalian subclades in red text possess nonclassical monocytes.

### Defining the nonclassical monocyte core regulatory complex

CoRCs are composed of transcription factors that allow cell types to enact phenotypic modules. Comparisons of CoRC activity and expression can therefore provide testable hypotheses of cellular inheritance between species. In order to evaluate the correlation of transcription factor activity with cell type identity in the monocyte compartment, we took advantage of the recently developed SCENIC workflow^42,43^. We chose to focus our efforts on a single-cell RNA-sequencing (scRNA-seq) dataset derived from human peripheral mononuclear blood cells (PBMCs)^37^. We first reanalyzed these data and manually assigned cell type identity, partially by referring to a previously reported gene-signature matrix^45^. Our investigation uncovered five major cell types differentiated by normalized gene expression in this compartment: classical dendritic cells (XCR1^hi^CD1C^hi^), plasmacytoid dendritic cells (CLEC4C^hi^IL3RA.1^hi^LILRA4^hi^), classical monocytes (CD14^hi^CD16^lo^), intermediate monocytes (CD14^+^CD16^+^), and nonclassical monocytes (CD14^lo^CD16^hi^) (**Supplementary Figure 1A**).

The SCENIC pipeline allows for the quantification of transcriptional network activity from scRNA-seq data by generating a correlation matrix between gene enrichment and transcription factor binding sites. When organizing these cells on a UMAP representing high-dimensional transcriptional-network space, we found that they clustered together by type (**Figure 2A**). We also observed polarity in the clustering of classical and nonclassical monocytes, with intermediate monocytes spread within the extremes of both cell types. Our findings suggest that mononuclear blood cell types can be classified according to their transcriptional network activity with high levels of accuracy. The transcription factors IRF8, KLF4, KLF2, C/EBPb, and NR4A1 compose a network required for the generation of nonclassical monocytes in humans and mice^7^. The expression of these proteins amongst the five major cell types was consistent with previous findings (**Supplementary Figure 1B**). KLF4 and IRF8 were expressed throughout the monocyte compartment, whereas C/EBPb was enriched in nonclassical monocytes (**Figure 2B; Supplementary Figure 1B**). NR4A1 and KLF2 were concentrated amongst both intermediate and nonclassical monocytes. We therefore decided to model the nCoRC using these five transcription factors (**Figure 2C**).

**Figure 2.**
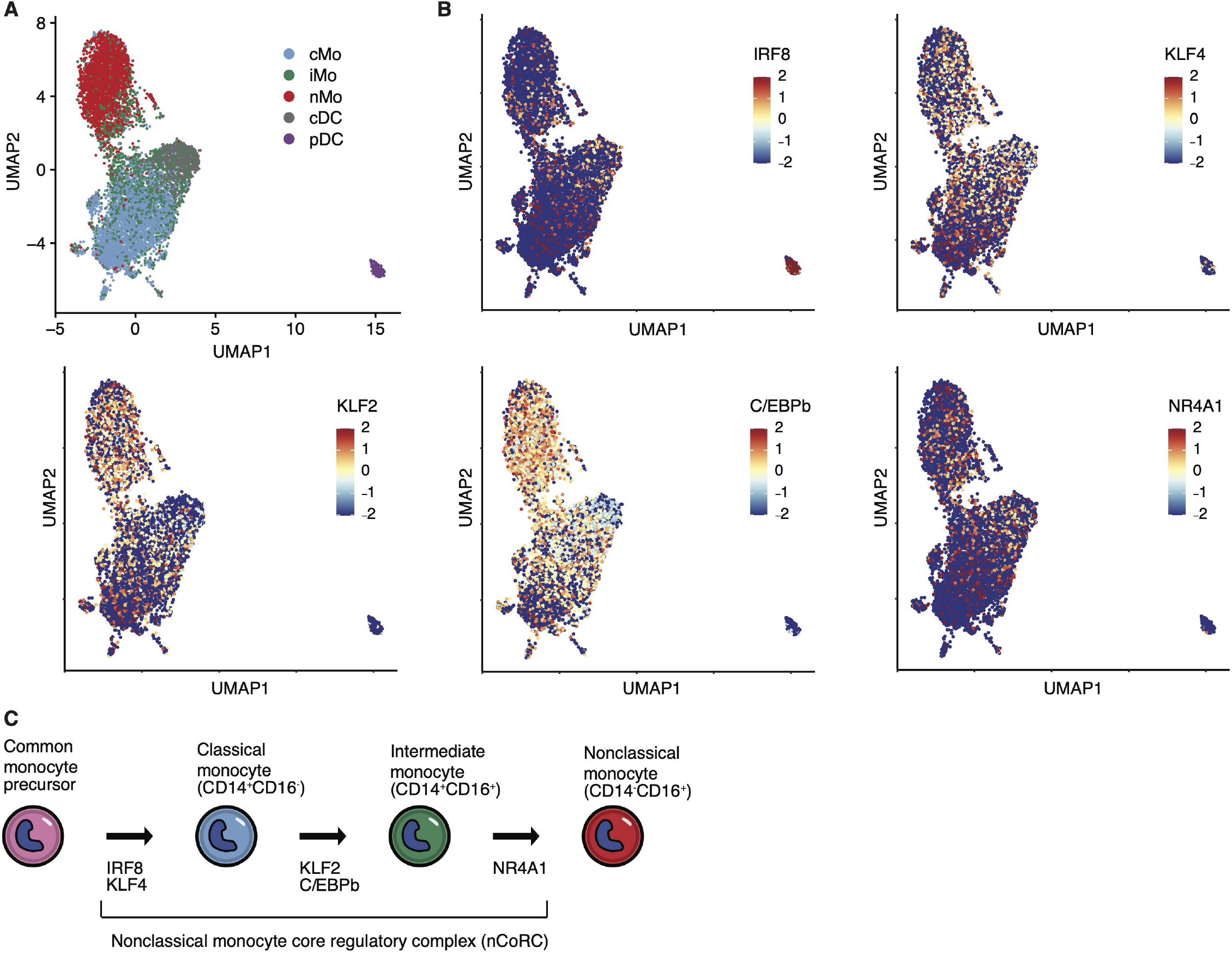
Regulatory network analysis of human monocytes highlights nCoRC proteins. (A) UMAP of human classical monocytes (cMo), intermediate monocytes (iMo), nonclassical monocytes (nMo), classical dendritic cells (cDC), and plasmacytoid dendritic cells (pDC) organization within high-dimensional transcriptional-network space. (B) Feature plots colored by the scaled expression of IRF8, KLF4, KLF2, C/EBPb, and NR4A1 amongst the five major cell clusters. (C) Diagram of nCoRC member activity during the conversion of classical monocytes into nonclassical monocytes.

### Members of the nCoRC are conserved amongst mammals

We next assessed the conservation of nCoRC members amongst major tetrapod groups through the use of a BLAST-based pipeline. We performed sequence similarity analyses on the transcription factors C/EBPb, IRF8, KLF2, KLF4, and NR4A1 amongst a selection of 68 tetrapod vertebrates with annotated genomes. These animals included amphibians, testudines, lepidosaurs, archosaurs, and mammals (**Figure 1; Supplementary Table 1**). The human MANE reference sequences for each of these proteins served as the baseline for proteome comparisons in a BLAST workflow. By averaging the similarity scores of these proteins between the human reference sequence and tetrapod clades, we found that the nCoRC exhibited 86% similarity amongst mammals (**Figure 3A**). This average level of homology was not common to archosaurs (61%), testudines (61%), lepidosaurs (65%), or amphibians (61%) (**Figure 3A**). The most conserved member of the nCoRC amongst mammals was NR4A1, with an average sequence similarity score of 91% (**Figure 3A, B**). NR4A1 expression is required for inducing the nonclassical monocyte program in both humans and mice^13^. The most conserved transcription factor amongst all clades was IRF8 at 79%. These data suggest that increased preservation of nCoRC homology is an attribute specific to the mammalian lineage. We then measured the variation in sequence similarities of individual nCoRC proteins within the mammalian clade and how this variation compares to aves (birds) and reptiles (crocodilians, lepidosaurs, and testudines). Median nCoRC sequence similarity was highest in mammalian proteins compared to those from either aves or reptiles (**Figure 3B**). Moreover, with the exception of KLF4, both aves and reptiles exhibited lower amounts of variation in sequence similarity of nCoRC proteins when compared to mammals (**Figure 3B**).

**Figure 3.**
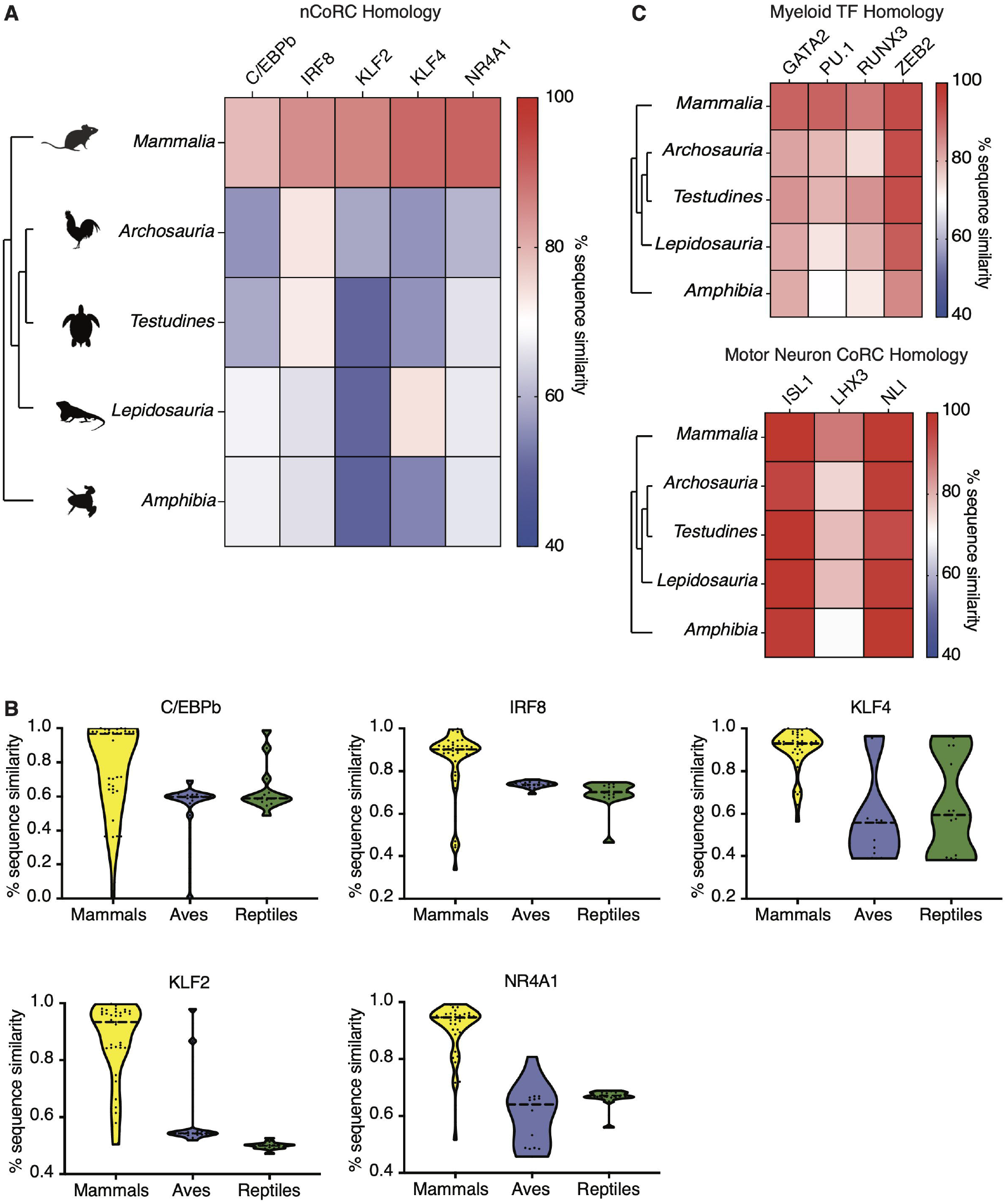
nCoRC protein sequence homology is conserved amongst mammals. (A) Heatmap of average nCoRC sequence similarity scores (% of human protein sequence) amongst major tetrapod clades. (B) Violin plots of homology scores between mammals, reptiles (crocodilians, lizards, and snakes), and aves. (C) Heatmaps of average sequence similarity scores for myeloid transcription factors (TFs; top) and the V2a spinal motor neuron CoRC (bottom) between major tetrapod clades.

Low levels of human nCoRC homology in non-mammalian versus mammalian tetrapods could be a general feature of all mammalian proteins. To test this alternative hypothesis, we assessed the conservation of myeloid transcriptional regulators GATA2, PU.1, RUNX3, and ZEB2^7^ between mammals, archosaurs, testudines, lepidosaurs, and amphibians. Unlike what we uncovered for nCoRC proteins, these transcription factors exhibited equal levels of sequence similarity between all clades considered (**Figure 3C**). As a negative control, we quantified the preservation of the CoRC proteins ISL1, LHX3, and NLI, which are responsible for the generation of V2a spinal motor neurons in amniotes^5^. We again found increased amounts of homology (> 88%) for these proteins amongst all tetrapods (**Figure 3C**). Our results suggest that the transcription factors that yield nonclassical monocytes are uniquely preserved amongst mammals and that this feature is not necessarily shared by other proteins associated with distinct cell types.

### Boreoeutherians drive mammalian nCoRC protein homology

Mammals comprise a diverse set of individual species and nCoRC preservation through this lineage could be explained by specific subfamilies. To address this issue, we asked whether nCoRC protein homology is conserved equally amongst placental mammals, as well as between placentals, marsupials, and monotremes. We determined that average nCoRC protein conservation was greatest in placental mammals (> 88%) (**Figure 4A, B**). Placental mammals that exhibited marked nCoRC conservation included the species of *Primata, Rodentia, Chiroptera, Cetartiodactyla*, and *Perissodactyla*, which are all boreoeutherian mammals (**Figure 4A**). Concordantly, the only boreoeutherian with reduced nCoRC sequence similarity was the single member of the clade *Scandentia.* While the sequence similarity scores of one *Atlantogenata* family member (*Echinops telfairi*) deviated from the scores of the other two species, this same individual contained the highest NR4A1 sequence similarity (**Figure 4B, C; data not shown**). Our results highlight the protein sequence diversity of transcription factors involved in nonclassical monocyte production within the mammalian clade.

**Figure 4.**
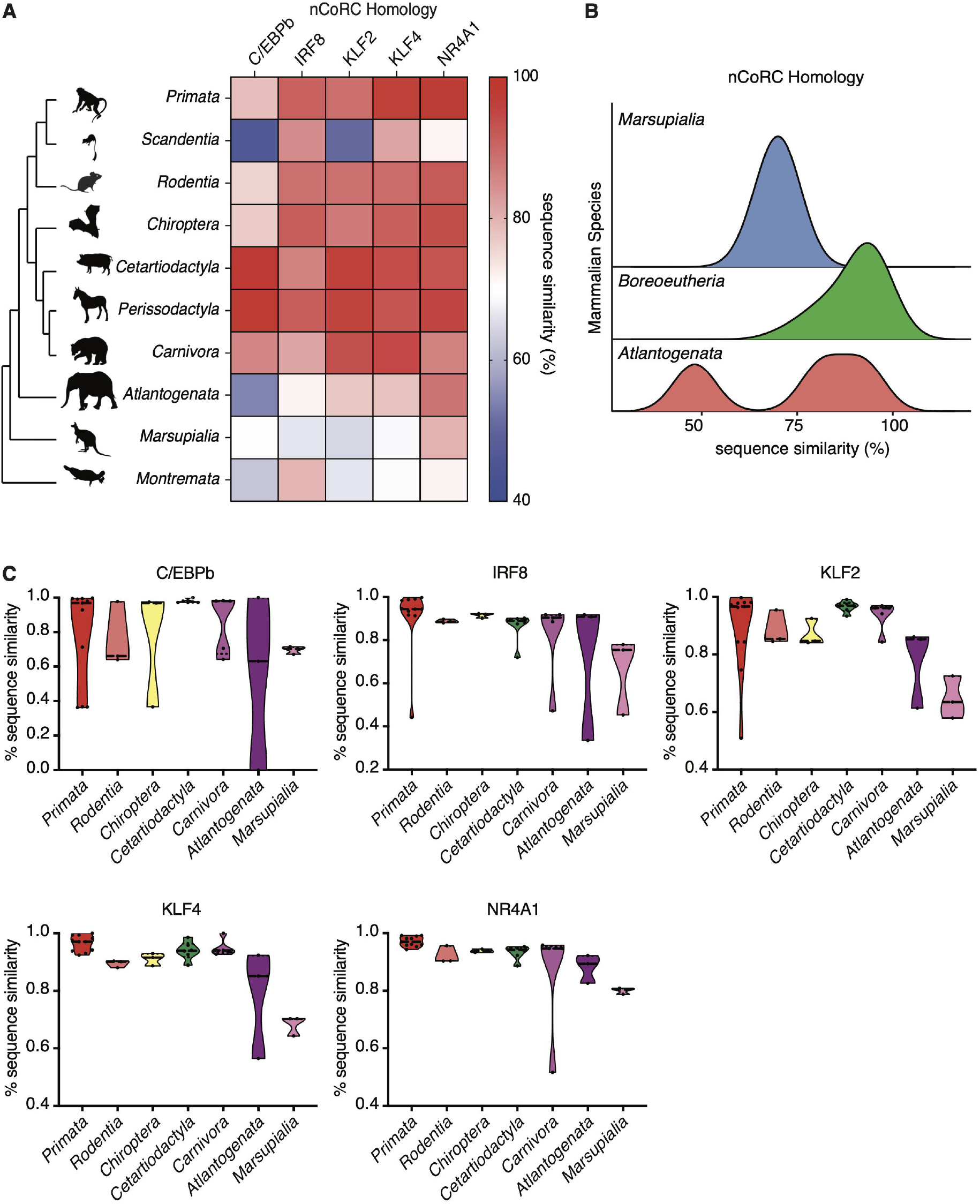
Conservation of mammalian nCoRC proteins is driven by boreoeutherians. (A) Heatmap of average nCoRC sequence similarity scores between placental mammals, marsupials, and monotremes. (B) Density plots of sequence similarity scores for individual species in the clades *Boreoeutheria, Atlantogenata*, and *Marsupialia.* (C) Violin plots of homology scores between *Atlantogenata, Marsupialia*, and subclades of *Boreoeutheria.*

We then sought to compare variations in the homology scores of individual nCoRC proteins between the subclades of *Boreoeutheria*, as well *Atlantogenata* and *Maruspialia.* C/EBPb and IRF8 displayed the greatest ranges of nCoRC sequence similarity amongst boreoeutherian clades (**Figure 4C**). We also noted that members of *Marsupialia* showed little variation in their nCoRC sequence similarity scores. (**Figure 4B, C**). These results highlight that while increased sequence similarity to human nCoRC proteins is a general feature of mammals, this effect is driven specifically by the members of *Boreoeutheria.*

### Receptors upstream of nonclassical monocyte production are conserved in mammals

Production of functional peripheral nonclassical monocytes is controlled by the activity of several receptors. The most famous of these include CSF1R, CX3CR1, NOTCH2, S1PR5, and TLR7. The activity of CSF1R and NOTCH2 induce the conversion of classical monocytes into nonclassical monocytes within the bone marrow^18,19^, with CSF1R expression being controlled by C/EBPb^46^. TLR7 acts independently and synergistically with NOTCH2 to increase nonclassical monocyte abundance^19^, whereas S1PR5 is necessary for nonclassical monocytes to exit the bone marrow^47^ and CX3CR1 is required for their survival in the periphery^48,49^.

We therefore quantified the sequence similarity scores of these proteins with the same BLAST workflow we used for nCoRC transcription factors. We determined that these receptor molecules were conserved by mammals at a level comparable to nCoRC proteins (> 81%; **Figure 5A**). This effect was again driven by boreoeutherian mammals (**Figure 5B**). Of these proteins, S1PR5 showed the greatest amount of variation within the mammalian clade. Our data indicate that the protein machinery responsible for nonclassical monocyte ontogeny has experienced marked levels of conservation on the mammalian lineage, presumably to preserve the ability of these species to generate nonclassical monocytes.

**Figure 5.**
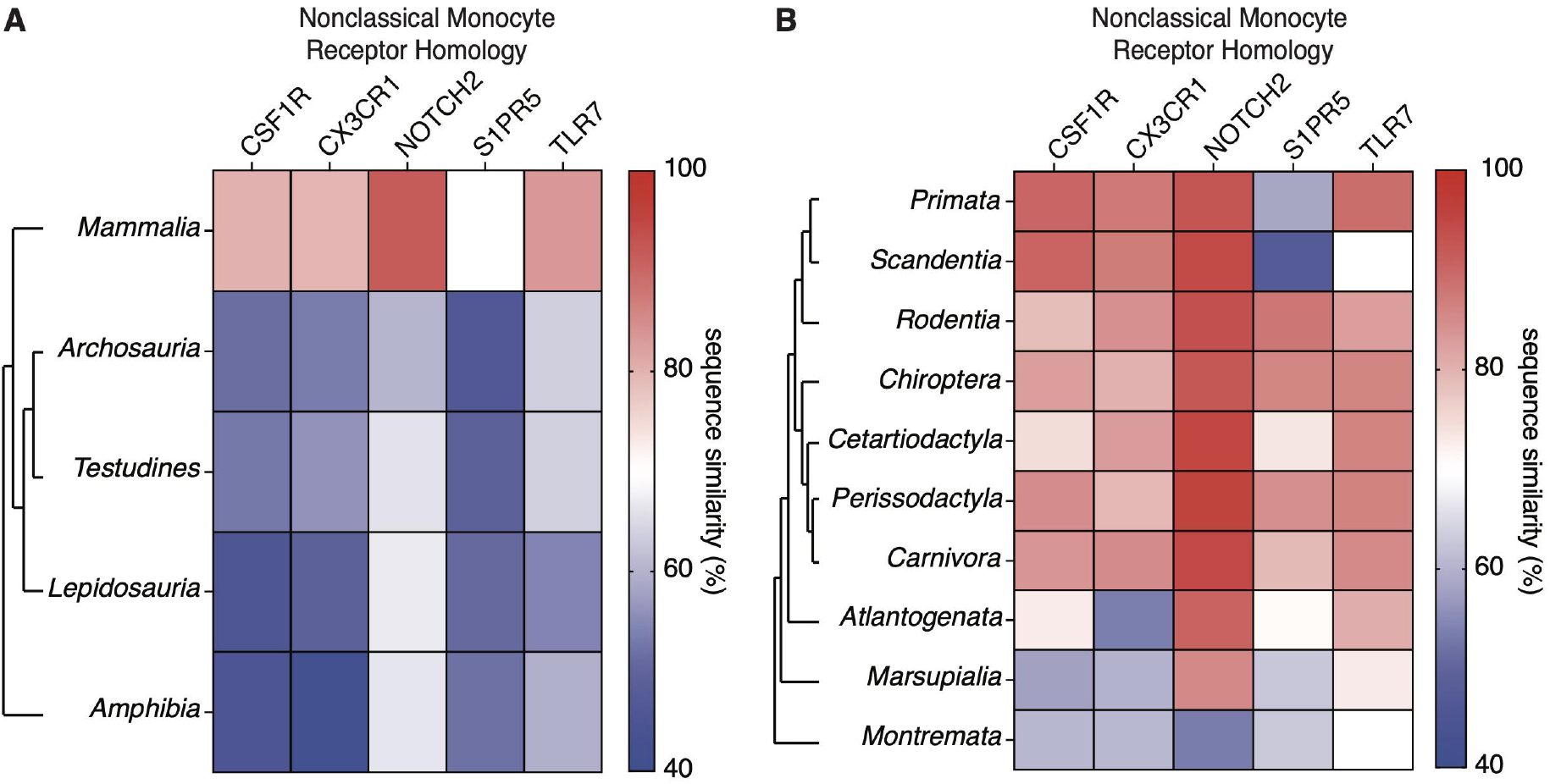
Receptors linked to nonclassical monocyte ontogeny are conserved in mammals. (A) Heatmap of average sequence similarity scores for receptors enriched on nonclassical monocytes between major tetrapod groups. (B) Heatmap of average sequence similarity scores for receptors enriched on nonclassical monocytes between placental mammals, marsupials and monotremes.

## DISCUSSION

Modeling the evolutionary history of immune cells remains challenging due to the lack of cellular remains in the fossil record and the diversity of observed cell types within extant animals. Nevertheless, recent advancements in cellular transcriptomics allow investigators to define cell types in terms of their transcriptional ontogeny and can provide methods for testing hypotheses of cellular ancestry^5,6^. Nonclassical monocytes are specialized, long-living monocytes that exit the bone marrow to crawl along and patrol the peripheral vasculature^14–17^. These cells have been found exclusively in peripheral blood and bone marrow of mammals. While monocytes are common to all tetrapods, whether nonclassical monocytes are restricted to mammals or if they are shared by other tetrapod clades has not been determined.

Herein, we analyzed a scRNA-seq dataset derived from healthy human PBMCs^37^ to define the CoRC of nonclassical monocytes. This analysis highlighted the expression of C/EBPb, IRF8, KLF2, KLF4, and NR4A1 within the monocyte compartment. We determined that these proteins are all uniquely conserved by boreoeutherian mammals and that this pattern is not observed for other myeloid transcription factors. KLF4 and IRF8 form a transcriptional circuit that biases common monocyte precursors to become classical monocytes^7,9^. Our laboratory has shown that KLF2 induces the nonclassical monocyte gene program through upregulation of NR4A1 expression^13^. C/EBPb also controls the expression of both NR4A1 and CSF1R by monocytes, and is required for nonclassical monocyte survival^11,46^. Thus, a more refined model of classical to nonclassical monocyte conversion is now emerging for the homeostatic bone marrow (**Figure 6**).

**Figure 6.**
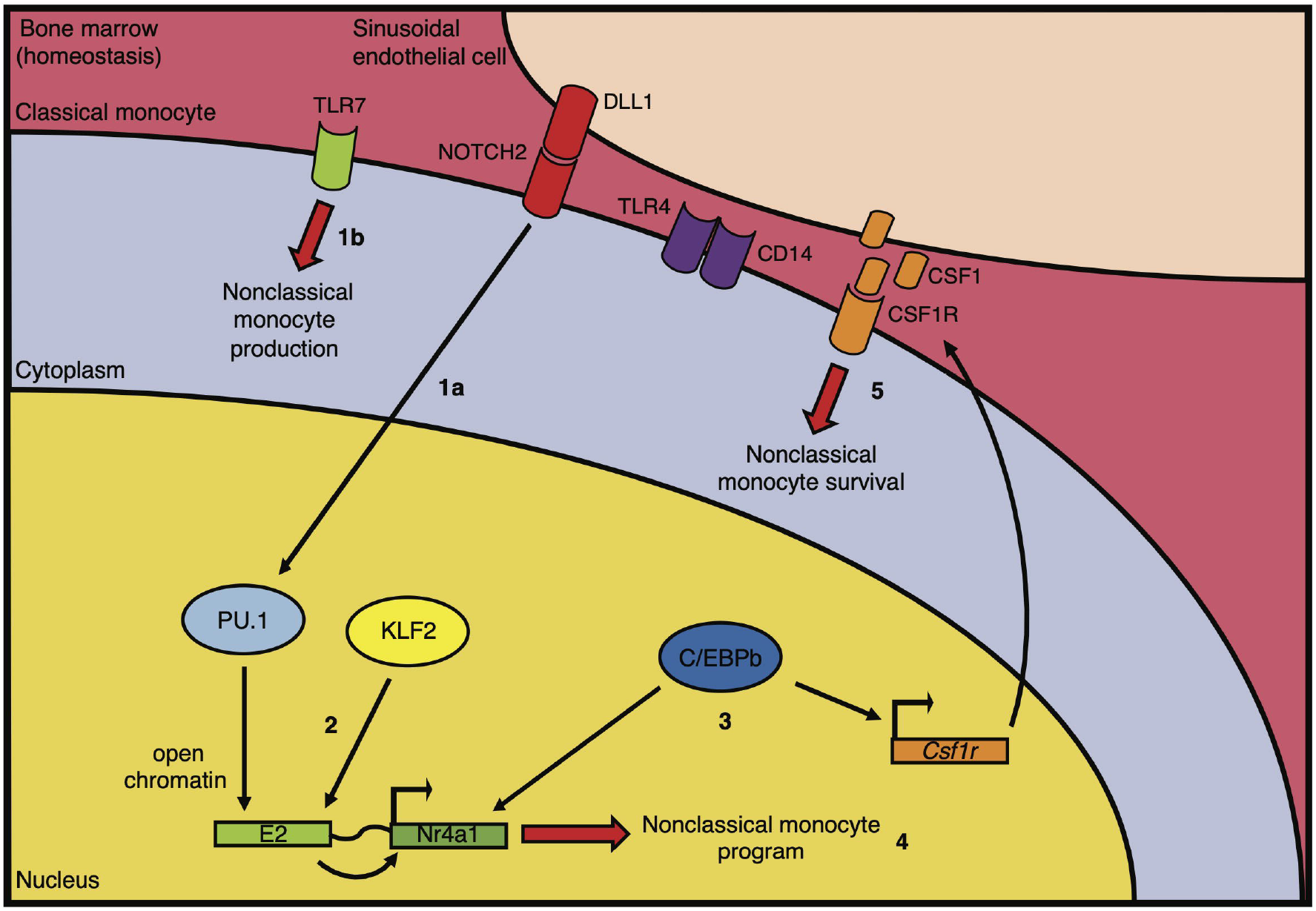
Model of classical to nonclassical monocyte conversion in the bone marrow. After the establishment of CD14^+^TLR4^+^ classical monocyte identity by the IRF8:KLF4 circuit within common monocyte precursors, (1a) NOTCH2 and (1b) TLR7 signaling independently and synergistically induce nonclassical monocyte production. Sinusoidal niches of NOTCH2^+^ endothelial cells cause PU.1 to open chromatin surrounding the E2 super-enhancer in CD14^+^TLR4^+^ monocytes. (2) KLF2 then upregulates the expression of NR4A1 through the E2 super-enhancer. (3) C/EBPb helps controls the expression of both NR4A1 and CSF1R. (4) NR4A1 expression subsequently induces the nonclassical monocyte program. (5) Release of CSF1 by sinusoidal endothelial cells promotes nonclassical monocyte survival.

The results of this report suggest that nonclassical monocytes are a synapomorphy of boreoeutherian mammals. Supporting this hypothesis is our laboratory’s observation that the nonclassical monocyte program driven by NR4A1 relies on the conserved E2 super-enhancer containing KLF2-binding motifs in both humans and mice^13^. Still, whether the E2 super-enhancer is common only to the family containing rodents and primates (*Euarchontoglires)* or if it is shared by a larger set of mammals is unknown. Birds have acquired both 4-chambered hearts and endothermic metabolism independently of mammals^50^. Thus, even though our protein homology results herein suggest otherwise, birds could still possess mononuclear cells that are either orthologous or homoplastic to mammalian nonclassical monocytes. Moreover, our hypothesis in which nonclassical monocyte ontogeny is an inherited trait of boreoeutherian mammals does not preclude derivations to this cell type within *Boreoeutheria.* For example, both classical (CD14^hi^CD163^lo^) and intermediate (CD14^lo^CD163^hi^) monocytes are present in pigs, but whether the latter population contains true nonclassical monocytes has not been answered^51^. Additional genomic profiling in combination with high-dimensional phenotyping and physiological examination of PBMCs, for both mammalian and non-mammalian species, will therefore be required to resolve the ancestry of nonclassical monocytes.

## ACKNOWLEDGEMENTS

This study was supported by P01 HL136275, U01 CA224766, and R01 HL134236. We thank the members of the Hedrick Laboratory for their helpful discussions and critiques of this work.

## AUTHOR CONTRIBUTIONS

D.J.A. conceptualized the study, performed data analysis, generated figures, and wrote the manuscript. D.F. performed BLAST-based protein analyses using tetrapod genomes, generated figures, and edited the manuscript. A.A. performed the SCENIC-based analyses of human monocyte scRNA-seq data, generated figures, and edited the manuscript. A.Z.N. contributed to the study design, performed data analysis, and edited the manuscript. C.C.H. oversaw the project and edited the manuscript.

---

**Supplementary Figure 1. Gene-expression profiling of human mononuclear blood cells**. (A) Hierarchical clustering of normalized (left) and scaled (right) immune cell marker expression amongst the five mononuclear blood cell types we identified in the scRNA-seq data of Watanabe et al., 2020. (B) Density plots of normalized IRF8, KLF4, KLF2, C/EBPb, and NR4A1 expression within the UMAP clusters constructed in **Figure 2A**.

**Supplementary Table 1. Descriptive characteristics of included tetrapod species.**

