## Supplementary figures and images for "Modeling the evolutionary history of nonclassical monocytes in mammals"

### Supplementary Figure 1

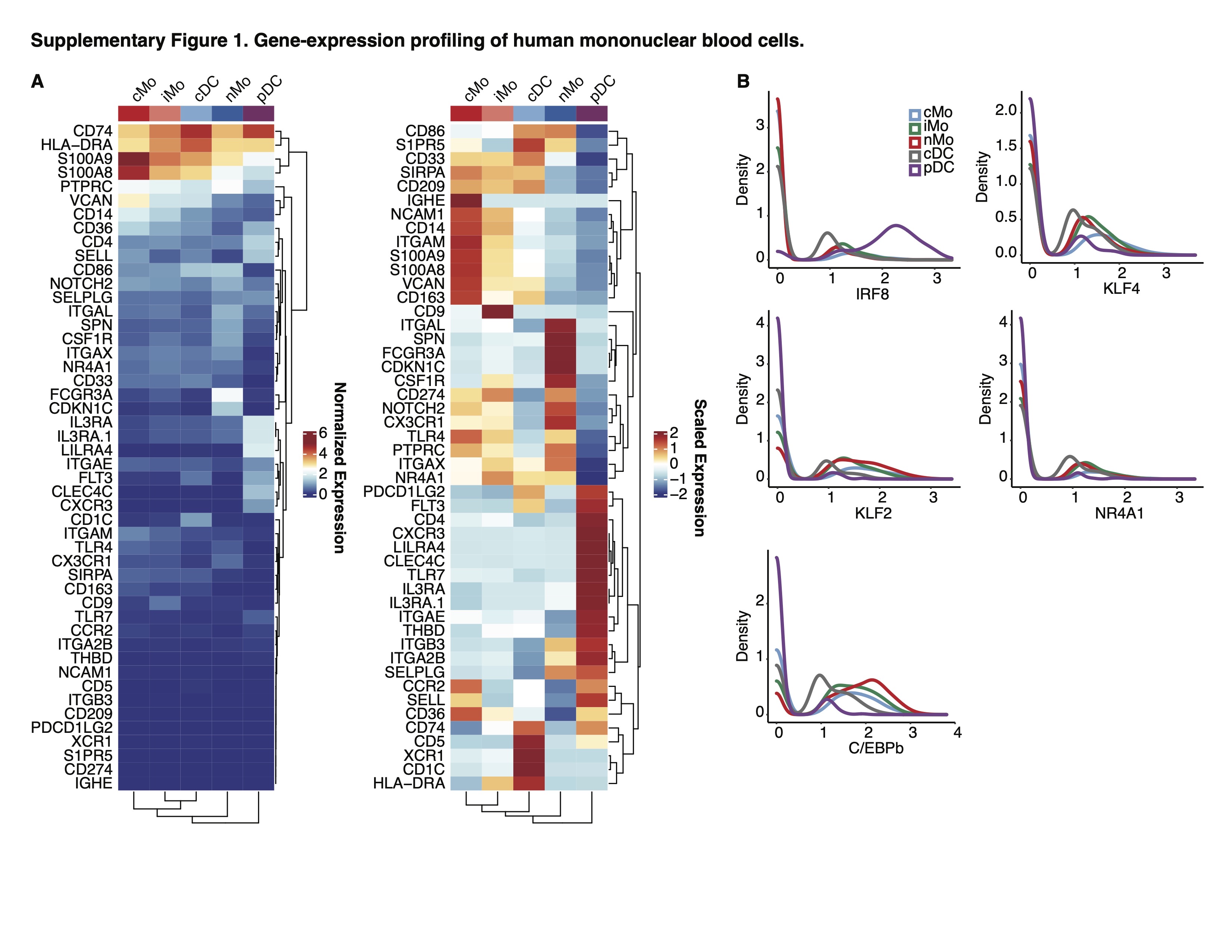
